# A conserved architectural domain shapes centromere evolution in *Drosophila*

**DOI:** 10.64898/2026.06.30.735607

**Authors:** Alejandra Samano, Mahul Chakraborty

## Abstract

Centromeres ensure faithful chromosome segregation despite being embedded within rapidly evolving repetitive DNA, a contradiction known as the centromere paradox. While centromere identity is defined by the histone variant CENP-A, how conserved function is maintained amid rapid DNA turnover remains unclear. Here, we generate highly contiguous genome assemblies from single *Drosophila melanogaster* individuals that, for the first time, resolve a chromosome through its centromere, linking the chromosome 3 arms within a continuous sequence. Comparative assemblies from wild-derived strains reveal extensive structural variation in pericentromeric satellites, including large-scale expansions, contractions, and sequence divergence. Despite this variation, the CENP-A–associated centromeric core exhibits conserved organization across strains. Integration of Hi-C interaction maps with sequence analyses shows that flanking *dodeca* satellite arrays form a spatially interacting domain that bridges both sides of the centromere, whereas adjacent *Prodsat* arrays are more variable and show weaker interactions. These results support a model in which rapidly evolving centromeric DNA is constrained by conserved higher-order architecture, providing a framework for reconciling the rapid evolution of centromere sequence with its conserved function.

## Introduction

Centromeres are essential chromosomal loci that mediate kinetochore attachment to spindle microtubules, ensuring accurate chromosome segregation during cell division. Despite this conserved function, centromeric DNA evolves rapidly across species, presenting a long-standing contradiction known as the “centromere paradox” (Henikoff et al. 2001; Kursel and Malik 2018). This paradox is thought to arise from centromere drive during asymmetric female meiosis and the compensatory evolution of kinetochore proteins, resulting in large, dynamic satellite arrays that shift over evolutionary time. One resolution to this paradox is that centromere identity is determined not by primary DNA sequence but by epigenetic and structural features that are maintained despite rapid sequence turnover. Although centromeres are epigenetically defined by the histone variant CENP-A, an additional layer of organization—such as higher-order chromatin structure or DNA topology—may provide a stable functional scaffold (Plohl et al. 2014; Kasinathan and Henikoff 2018). Under this model, repeat composition can change rapidly while preserving a conserved architectural context required for centromere function. However, how these constraints are maintained despite extensive sequence turnover remains unclear.

Addressing this question requires delineating the relationship between the evolutionary dynamics of the centromere–pericentromere sequences and their three-dimensional organization. Comparative analyses of highly contiguous assemblies in humans and plants have recently provided nucleotide-resolution maps of centromere structure and variation, leading to inferences of the organization and dynamics of repeats in these regions (Naish et al. 2021; Nurk et al. 2022; Logsdon et al. 2024; Xie et al. 2025). However, these studies have largely emphasized the linear organization, sequence turnover, and epigenetic state of centromeric repeats, leaving the relationship between three-dimensional architecture and the evolutionary dynamics of centromeric DNA less well understood. To address this gap, we use single-fly DNA to de novo assemble the genomes of three *Drosophila melanogaster* strains, enabling recovery of centromere and pericentromere sequences that have remained fragmented in previous assemblies (Chang et al. 2019). By integrating these assemblies with CENP-A profiles and high-resolution 3D interaction maps, we investigate how spatial organization relates to sequence variation across centromeric regions. Together, these data suggest that a subset of centromere-associated satellites exhibits hallmarks of evolutionary constraint, potentially linked to their incorporation into a conserved spatial domain

## Results

Long-read-based de novo genome assemblies of *D. melanogaster* recover fragments of centromeric islands, partly because genetic variation complicates assembly of centromeric and pericentromeric sequences (Courret and Larracuente 2023). To uncover the complete sequence of a *Drosophila* centromere-pericentromere region, we *de novo* assembled the genome of a single female of the near-isogenic *D. melanogaster* strain A4 (Chakraborty et al. 2018) using high coverage (68×) PacBio HiFi reads. The resulting assembly is highly contiguous (contig N50 = 24.5 Mb), accurate at the nucleotide level (QV = 53.6), and comparable to the reference genome in gene completeness (99.7% Dipteran BUSCOs)(Table 1, Supplemental Fig. 1)(Adams et al. 2020; Jia et al. 2024; Shukla et al. 2025). The A4 single fly assembly resolves repetitive regions that have remained fragmented in previous *D. melanogaster* genomes, including large pericentromeric satellite arrays (Shukla et al. 2025; Liu et al. 2025) (Fig. 1A, Supplemental Fig. 2). Notably, we recover a single 8.84 Mb contig that joins the arms 3L and 3R across the centromere, whereas the arms remain separated in existing assemblies (Fig. 1A, B; Fig. 2A). This large-scale organization is further supported by Hi-C contact maps, which show chromosome-wide interaction patterns consistent with correct scaffolding of the chromosome arms (Fig. 1C). This continuity provides a direct framework for examining centromere organization and the surrounding pericentromeric landscape within a single contiguous sequence.

**Figure 1.**
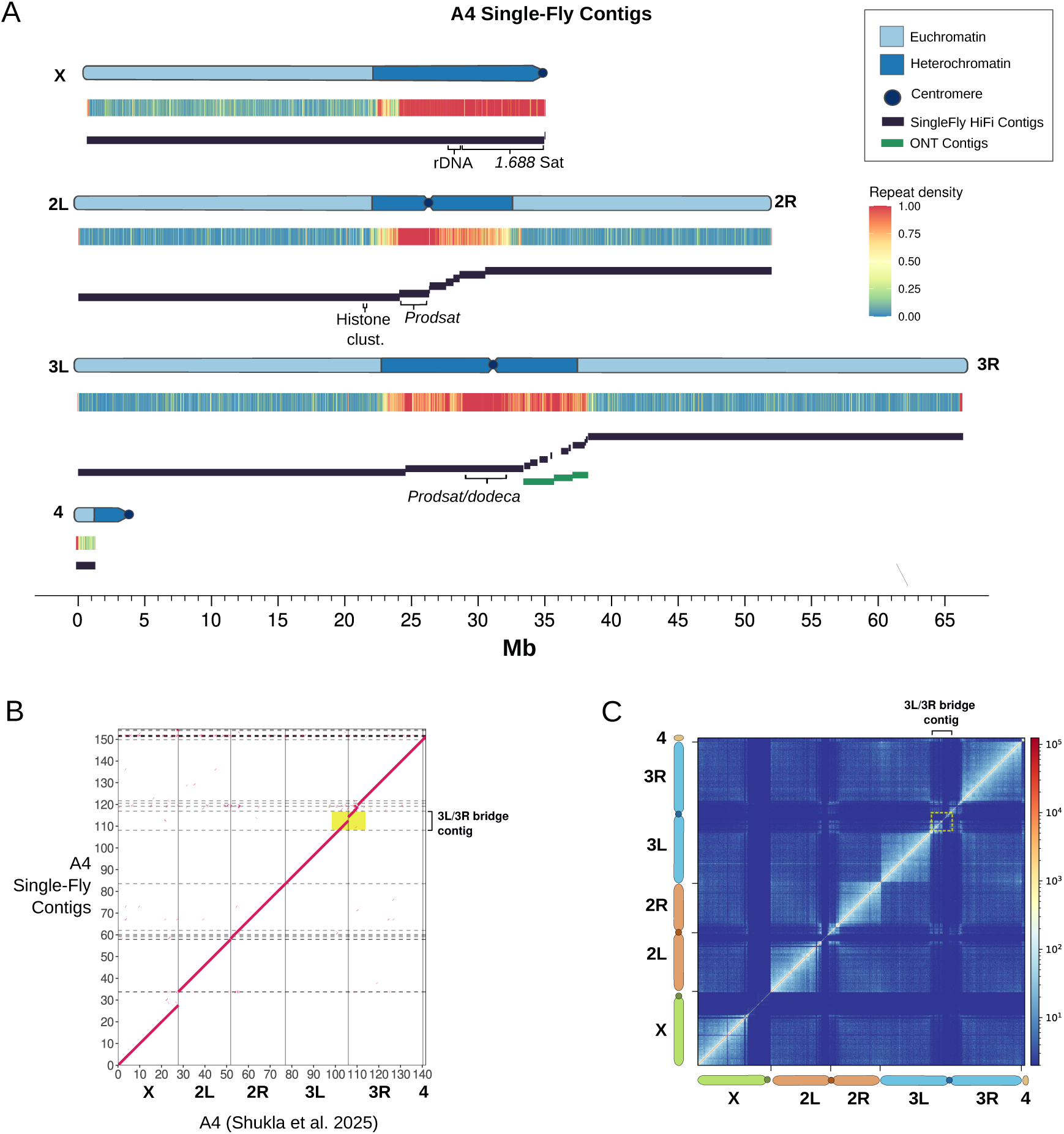
High-contiguity genome assembly of a single *Drosophila melanogaster* (A4) female. A. Chromosomal organization and repetitive landscape. Linear map of assembled contigs (black bars) aligned to major chromosome arms, showing high contiguity across both euchromatin and pericentromeric heterochromatin. Heatmaps indicate repeat density, with annotations for complex repetitive loci captured within single contigs, including the rDNA, 1.688 satellite, and *Prodsat*/*dodeca* arrays. B. Dot plot comparison between the previously published A4 HiFi-based assembly (x-axis) and the single-fly assembly (y-axis), showing high concordance across major chromosome arms with minimal structural discordance. The single-fly assembly recovers a contig spanning the 3L and 3R arms (yellow box). C. Hi-C contact matrix of the A4 single-fly assembly scaffolded into the major chromosome arms. The 3L/3R bridge contig (yellow dashed box) connects the 3L and 3R arms into a ∼65 Mb chromosome 3 scaffold.

**Figure 2.**
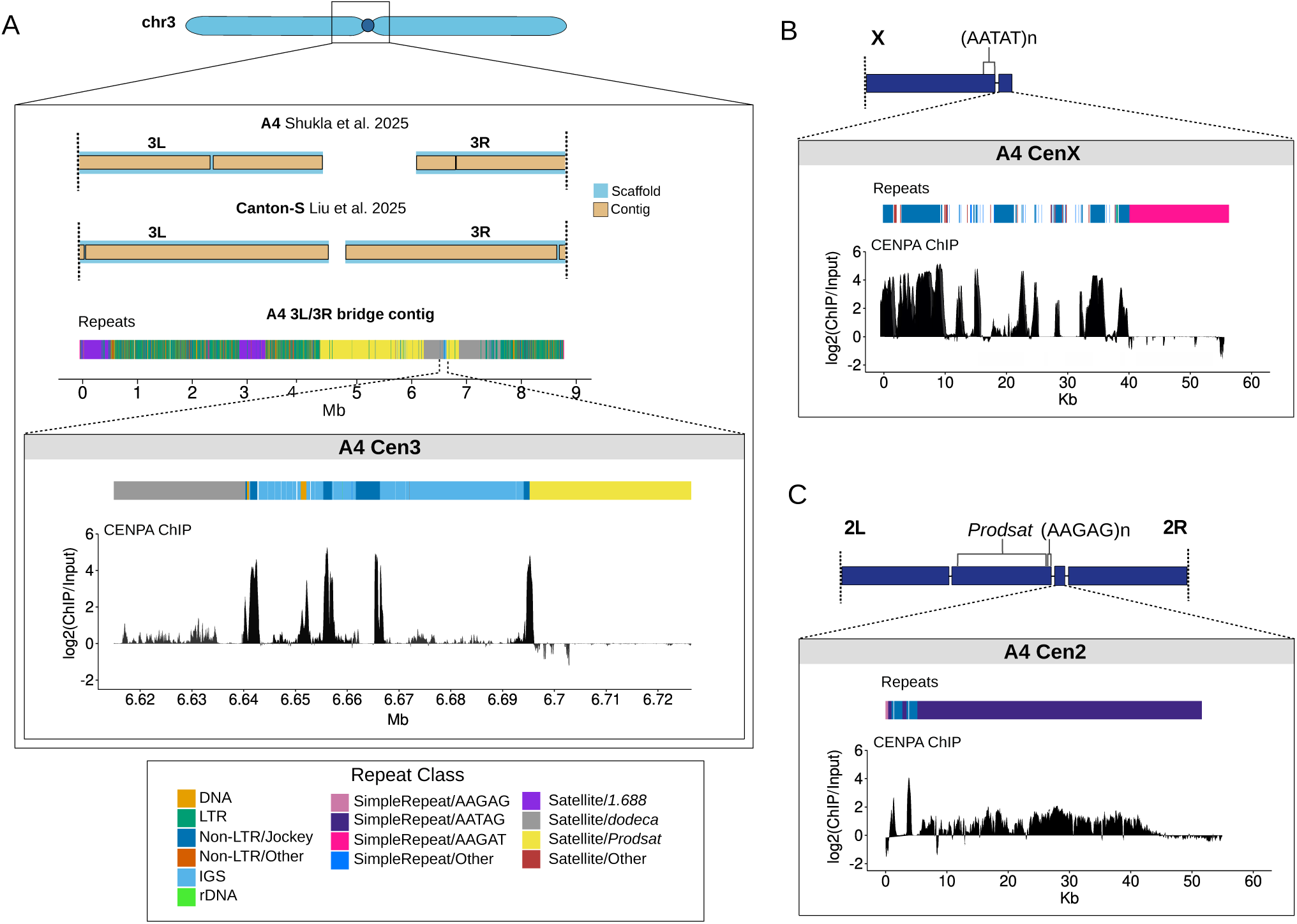
Contiguous assembly and CENP-A validation of *Drosophila melanogaster* centromeres. A. Top: Genomic alignment comparing the previous A4 assembly (Shukla et al. 2025) and the recently published Canton-S assembly (Liu et al. 2025), in which the centromeric region is fragmented across separate 3L and 3R scaffolds, with the A4 single-fly assembly, which resolves the region within a single ∼9 Mb contig. Bottom: Detailed view of the A4 Cen3 locus showing the repeat annotation and CENP-A ChIP-seq enrichment. B. Repeat annotation and CENP-A occupancy of the A4 CenX contig, with a schematic showing the placement relative to the X chromosome contig, which ends in AATAT repeats. C. Repeat annotation and CENP-A occupancy of the A4 Cen2 contig, including its placement relative to the 2L and 2R contigs, as well as the *Prodsat*-rich contig linked to 2L.

**Table 1.**
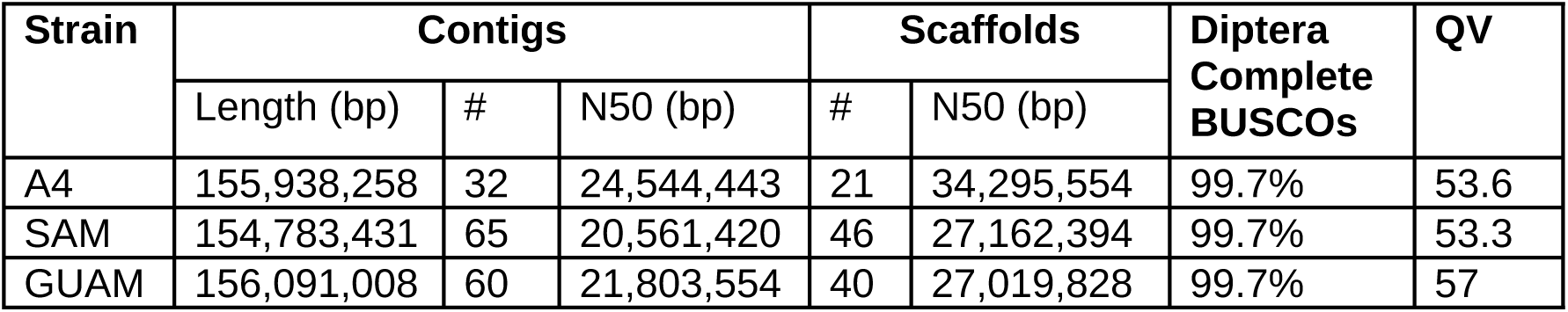
Primary assembly quality and completeness.

Mapping of raw HiFi reads to the assembly reveals uniform coverage across euchromatin, with reduced coverage across simple satellite arrays and pericentric heterochromatin, consistent with known sequencing biases and mapping ambiguities in low-complexity, highly repetitive DNA (Supplemental Figs. 3, 4) (Nurk et al. 2022; Xie et al. 2025; Shukla et al. 2025). Long-read Oxford Nanopore sequencing further supports the assembly and provides coverage across AT-rich satellite blocks that are underrepresented in HiFi data, highlighting platform-specific sequencing biases and offering complementary recovery of these regions (Supplemental Fig. 5). In addition to chromosome 3, we recover substantial portions of the centromere-associated sequences of chromosomes X and 2 (Fig. 1A). Together, these results show that single-fly assemblies recover centromeric and pericentromeric regions with sufficient continuity for detailed analysis of their sequence organization and structural context.

### Reconstruction of a contiguous chromosome

We defined the centromeric region in our assemblies based on previously described retroelement-rich architecture and CENP-A enrichment patterns (Chang et al. 2019). The chromosome 3 centromeric contig shows significant CENP-A enrichment peaks primarily localized to the retroelement-rich ’islands’ (Chang et al. 2019)(Fig. 2A). The organization of this region is consistent with the previously described architecture of the chromosome 3 centromere, with *G2/Jockey-3* elements exhibiting high CENP-A occupancy, separated by the intergenic spacer of rDNA (IGS) sequences and flanked by *Prodsat* (the 10 bp satellite) and *dodeca* satellites (Chang et al. 2019). These observations indicate that the assembly spans the centromeric region and links the chromosome 3 arms within a continuous sequence.

We also recovered contiguous blocks containing the centromere-associated sequences of chromosomes X and 2. Although these sequences are not physically joined to their respective chromosomal arms – likely due to the large, near-identical 5-bp simple repeat arrays – their recovery enables analysis of centromere sequence organization and repeat composition across multiple chromosomes. The X-centromere contig (CenX; 53.6 Kb) is concordant with previous characterizations of this region, including those derived from the *Dp1187* minichromosome, long-read sequencing, and fluorescence in situ hybridization (FISH) (Sun et al. 2003; Chang et al. 2019), and exhibits CENP-A enrichment over *G2/Jockey-3* elements flanked by AAGAT repeats (Fig. 2B). Similarly, the chromosome 2 centromere contig (Cen2; 93.3 Kb) consists of a small ∼4 Kb core containing two *G2/Jockey-3* elements with strong CENP-A enrichment embedded within AATAG repeats (Fig. 2C).

Our X chromosome contig recovers ∼10.7 Mb of additional pericentromeric sequence relative to the iso-1 release 6 reference assembly (Hoskins et al. 2015), including the rDNA array, where the most recent HiFi-based A4 assembly terminates (Shukla et al. 2025) (Fig. 1B, Supplemental Fig. 6). This extension contains a ∼6 Mb array of 1.688 satellite DNA and ends with AATAT repeats. As these repeats are known to immediately flank the centromeric region (Sun et al. 1997; Sun et al. 2003), this suggests that our contig extends well into the centromere-proximal satellite arrays.

While the functional core of *D. melanogaster* centromeres has been characterized, resolving the large satellite arrays flanking them has remained a major challenge in heterochromatin assembly. The pericentric satellite content in our A4 assembly is consistent with experimental estimates. For example, the 12-bp *dodeca* satellite, which is highly concentrated in arrays flanking Cen3, is estimated to comprise approximately 1 Mb of the genome (Abad et al. 1992), and our assembly recovers 993,068 bp. *Prodsat* is a major constituent of the pericentromeres of chromosomes 2 and 3 and typically ranges from 2 to 5 Mb (Lohe et al. 1993; Török et al. 2000; Wei et al. 2014; Talbert et al. 2018); our A4 assembly includes 4.52 Mb of *Prodsat* sequence. While the *Prodsat* and *dodeca* arrays flanking Cen3 are incorporated within the arm-joining contig (Fig. 2A), the pericentromeric satellites of chromosome 2 remain partially fragmented. We identified a pericentromeric 2.2 Mb contig based on its contact frequency with 2L. This contig is composed almost entirely of *Prodsat*, interspersed with retroelements and bounded by simple 5-bp repeats, consistent with previous estimates of the chromosome 2 *Prodsat* array. (Supplemental Fig. 7, Supplemental Table 4) (Török et al. 2000).

### Assembly of Wild-Derived Strains

To examine genetic variation in centromeric and pericentromeric regions across diverse genetic backgrounds, we *de novo* assembled the genomes of two wild-derived strains from Guam (GUM) and American Samoa (SAM) using DNA from single females (Supplemental Table 1). These assemblies are comparable to the A4 assembly in contiguity and completeness (Table 1) (contig N50: GUM 21.8 Mb; SAM 20.6 Mb). In contrast to the highly inbred A4 strain, the GUM and SAM assemblies resolved two distinct haplotypes, revealing substantial structural heterogeneity, including several large inversions (Supplemental Tables 2, 3). In GUM, we recovered both the standard and inverted arrangements of *In(2L)t*, but only the *In(3R)Payne* arrangement (Corbett-Detig and Hartl 2012)(Supplemental Fig. 8). Mapping of raw HiFi reads back to the GUM assembly shows no split reads at the inversion breakpoints, suggesting that the sequenced individual was homozygous for *In(3R)Payne* (Supplemental Fig. 9). In SAM, both the standard and inverted arrangements of *In(3R)Payne* were recovered, along with the common *In(2R)NS* inversion. In addition, SAM harbors a 7.7 Mb inversion on chromosome 2L that, to our knowledge, has not been previously described and may represent a rare or population-specific variant (Supplemental Fig. 10). These results demonstrate that single-fly assemblies capture large structural variants at the chromosome scale, consistent with known patterns of inversion polymorphism in *D. melanogaster* populations.

Importantly, assemblies of both strains reconstruct a single, gapless sequence spanning the chromosome 3 centromere, uniting the 3L and 3R arms as in A4. Mapping CENP-A ChIP-seq data (Chang et al. 2019) to the GUM and SAM assemblies shows strong enrichment peaks within these arm-spanning contigs, as well as in unplaced contigs with sequence organization similar to the A4 centromere-associated regions of chromosomes X and 2 (Supplemental Fig. 11). The consistent recovery of the chromosome 3 bridge and centromere core contigs across distinct genetic backgrounds, together with the detection of inversion polymorphisms, supports the robustness of these assemblies and enables direct comparison of centromeric organization and pericentromeric architecture across *D. melanogaster* strains.

### Comparative analysis of functional centromeres and pericentric architecture

Using these assemblies, we next compared the structure of the retroelement-rich CENP-A–associated regions and surrounding pericentromeric sequences across strains. Comparative structural analysis of the retroelement-rich CENP-A domains (excluding the flanking satellites) across our three assemblies and the iso-1 reference (Chang et al. 2019) revealed a broadly conserved organization, with discrete strain-specific differences. The Cen3 hub exhibits similar repeat composition across strains, with a core architecture consisting of *G2/Jockey-3* elements interspersed with IGS sequences and fragments of DNA transposons (Fig. 3A). Notably, several structural features appear conserved within the species: the leftmost *G2/Jockey-3* element is consistently fragmented and interrupted by a *Transib5* DNA element, and a *Protop A* DNA element is present in all assemblies and co-localizes with a peak of CENP-A enrichment. Sequence identity analysis of aligned blocks indicates that, with the exception of IGS unit insertions and a single *G2/Jockey-3*, A4 Cen3 is nearly identical to iso-1, suggesting these strains share a recent centromeric ancestor. The high degree of structural similarity suggests that Cen3 architecture, comprising the *G2/Jockey-3* and the DNA elements, is evolutionarily constrained.

**Figure 3.**
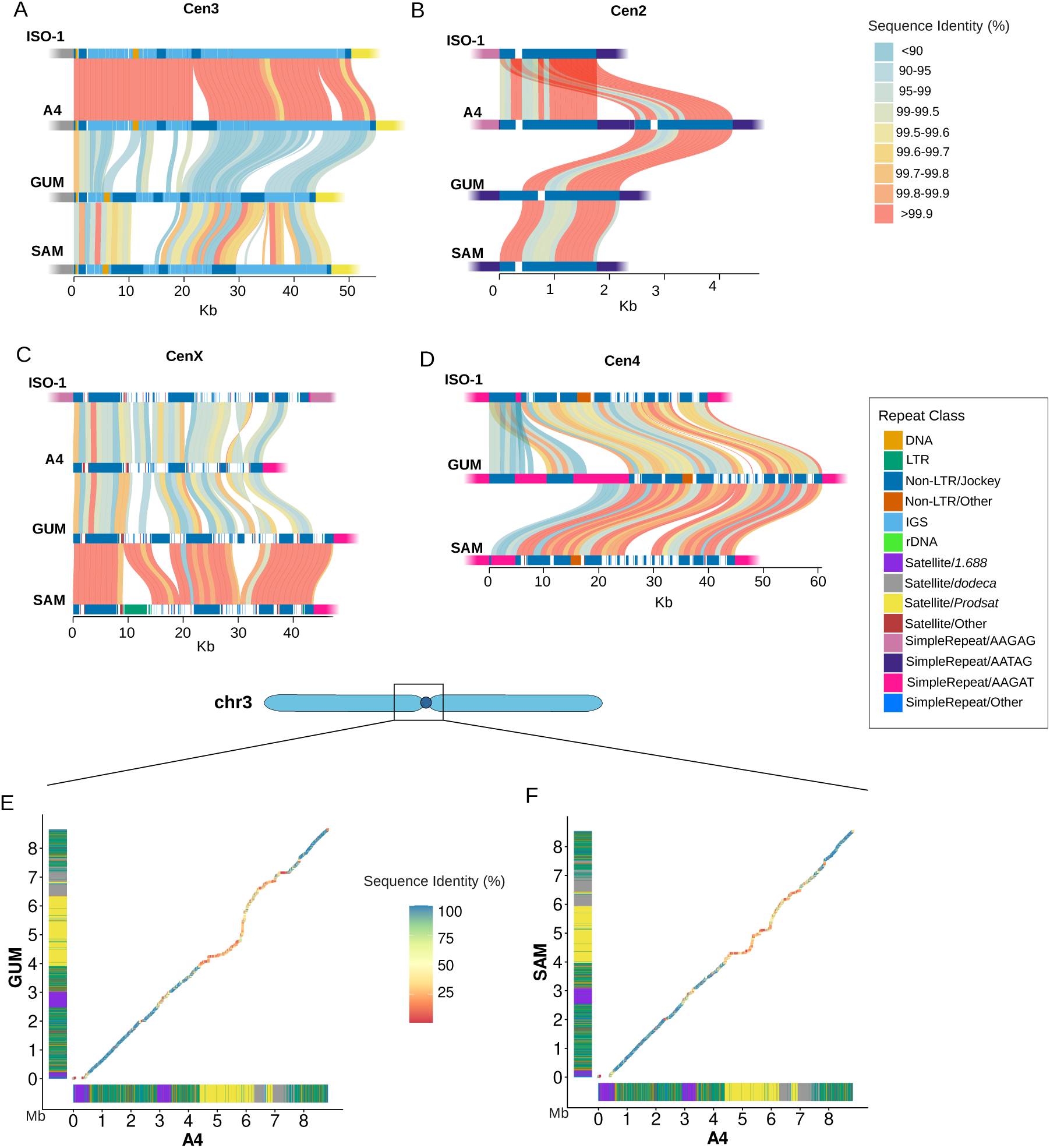
Comparative structural architecture and sequence divergence of *D. melanogaster* centromeric regions across strains. Alignments of the retroelement-rich CENP-A domains across the reference strain iso-1 (Chang et al. 2019) and three newly assembled strains are shown for (A) Cen3, (B) Cen2, (C) CenX, and (D) Cen4. Syntenic blocks between assemblies are colored by sequence identity (1kb windows). Flanking satellite blocks are excluded from the alignment and indicated by a gradient fade. (E–F) Pairwise alignment dot plots comparing the A4 chromosome 3 bridge contig against the corresponding assemblies from (E) GUM and (F) SAM.

In contrast to the stability of Cen3, the hubs of Cen2 and CenX exhibit more structural rearrangements (Figs. 3B, C). While GUM and SAM Cen2 maintain the single *G2/Jockey-3* motif as iso-1, the A4 strain harbors a duplication of this sequence and some of the adjacent AATAG repeat sequence. CenX remains largely syntenic across most strains; however, the SAM CenX is uniquely interrupted by a 4.5 kb *micropia* LTR insertion (Fig. 3C), consistent with the ongoing retrotransposon activity in centromeres (Wong and Choo 2004). In addition, the A4 CenX contains a strain-specific 3 Kb inversion encompassing IGS and non-repetitive sequence (Fig. 3C), further differentiating its architecture from the other strains.

We recovered a sequence similar to the Cen4 described in Chang et al. in our GUM and SAM assemblies, but it was absent from A4. Mapping A4 single-fly HiFi or Nanopore reads to the iso-1 Cen4 reference showed no aligned reads (Supplemental Fig. 12), and this lack of coverage was not observed in SAM or GUM. Consistent with this, the Cen4 sequence is also absent from previously published HiFi- and CLR-based A4 assemblies (Chakraborty et al. 2018; Shukla et al. 2025). As we identified no alternative regions of CENP-A enrichment in our A4 assembly, this suggests that the A4 Cen4 may be highly divergent relative to iso-1, potentially limiting detection using existing CENP-A datasets derived from a different genetic background (Chang et al. 2019).

### Structural and sequence variation across pericentric heterochromatin

Beyond the functional CENP-A hubs, we examined the pericentric heterochromatin of chromosome 3. Chromosome-wide comparisons revealed that this pericentromeric interval is the most divergent portion of the chromosome 3 scaffold in both GUM-A4 and SAM-A4 comparisons (Supplemental Fig. 13). Within this region, comparative alignments of the chromosome 3 bridge contigs show that sequence conservation is highest within TE-rich blocks (Figs. 3E, F). This pattern is likely driven by the high concentration of LTR retrotransposons, which, as relatively recent additions to the *D. melanogaster* lineage (Pianezza et al. 2025), retain greater sequence homogeneity than the surrounding satellite DNA. In contrast, the satellite arrays remain highly dynamic; both GUM and SAM exhibit localized expansions of the *Prodsat* and *dodeca* arrays relative to A4, consistent with the rapid turnover expected in these repetitive regions (Figs. 3E, F). Despite this variation, the overall organization of the chromosome 3 pericentromere is conserved across strains, with the CENP-A–associated hub flanked by *Prodsat* and *dodeca* arrays. Self-alignment sequence identity heatmaps reveal substantial heterogeneity in sequence identity within these arrays. The extensive *Prodsat* array on the 3L side of the centromere spans ∼2 Mb and contains distinct blocks of high sequence identity (>95%) interspersed within broader regions of lower identity (∼90%). Although the overall size of this *Prodsat* array is broadly similar across strains (A4: 2.07 Mb, GUM: 2.3 Mb, SAM: 2.02 Mb), SAM and GUM contain localized high-identity regions within the array, consistent with recent homogenization or turnover. Notably, these high-identity regions are frequently associated with repeats also present in the Cen3 CENP-A domain. In SAM, one such region lies adjacent to a cluster of *G2/Jockey-3*, *gypsy* LTR, and IGS sequences, whereas in GUM, similar clusters occur in two high-identity regions. This pattern suggests that TE-rich sequences resembling Cen3 can become embedded within expanding *Prodsat* arrays and propagate along with surrounding repeat DNA. In contrast, the *dodeca* arrays, which are located closer to the CENP-A–associated hub, show lower overall sequence identity (85–90%) and less evidence of recent homogenization.

Despite extensive internal sequence heterogeneity and rapid satellite turnover, the contrast between rapid sequence turnover and conserved spatial organization highlights a separation between sequence evolution and higher-order chromatin structure in centromeric regions. Hi-C interaction maps of the A4 and SAM pericentromeres on chromosome 3 reveal strongly concordant interaction profiles between the two strains (Pearson r=0.872, Spearman ρ=0.807, *p*-value=0.024), indicating conservation of broad interaction organization despite local variation. We next examined whether this conserved organization was associated with specific satellite classes. In both strains, *dodeca* arrays flanking the CENP-A-enriched hub show strong local and long-range contacts, including interactions between arrays on opposite sides of the hub (Figs. 4A, B). To quantify this pattern, we classified 25-kb Hi-C bins by their dominant satellite content and modeled raw contact counts between satellite-pair classes. In A4, *dodeca–dodeca* contacts were 2.4-fold higher than *dodeca–Prodsat* contacts and 2.7-fold higher than *Prodsat–Prodsa*t contacts (*p*-values: 4.8×10^-9^ and <1×10^-10^). In SAM, the corresponding enrichments were 1.7-fold and 2.4-fold (*p*-values: 1.4×10^-5^ and <1×10^-10^). Together, these results suggest that *dodeca* does not simply occupy linear positions adjacent to the centromere but instead forms a spatially interacting domain surrounding the CENP-A hub.

**Figure 4.**
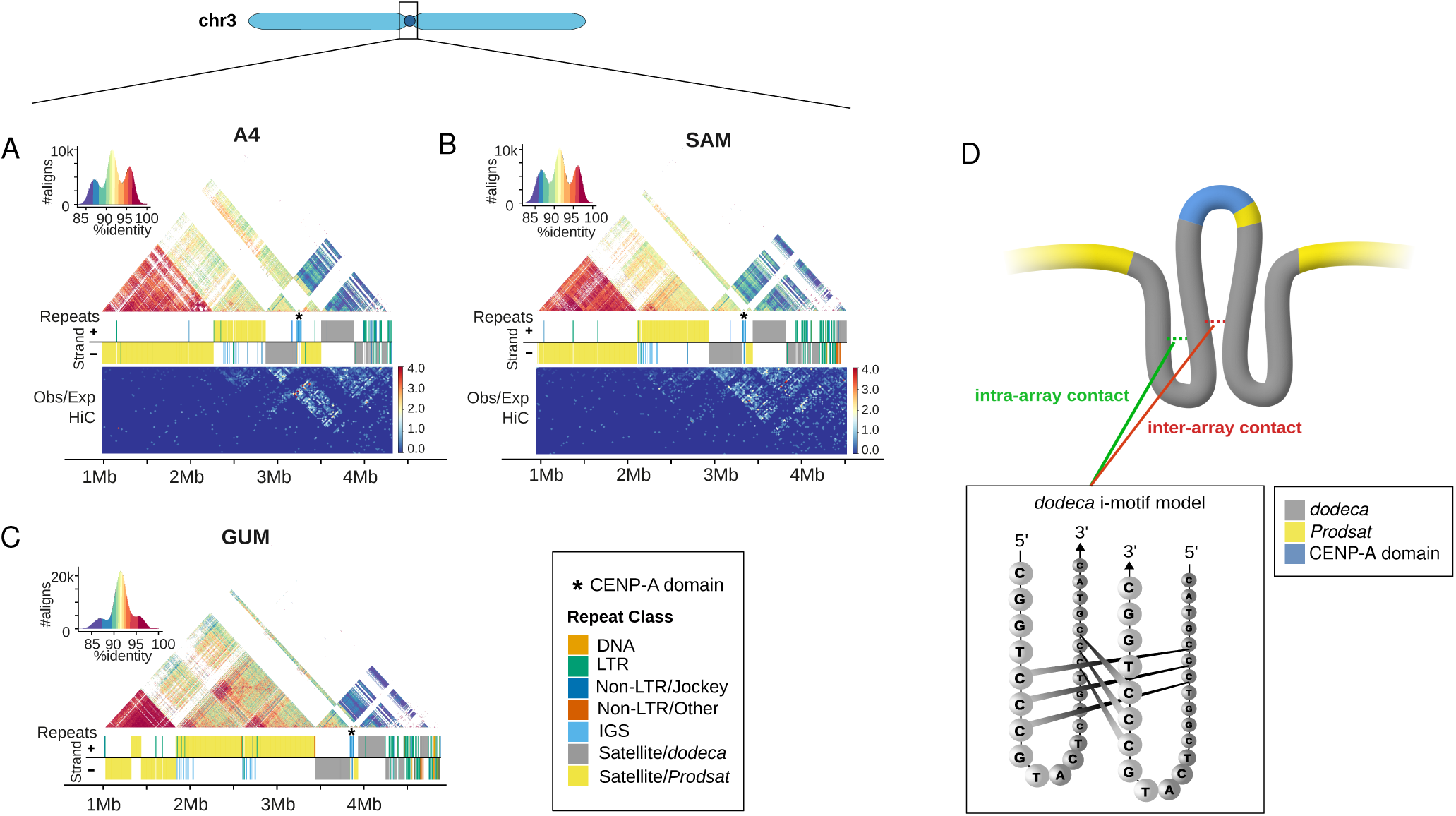
Sequence organization and three-dimensional interactions of chromosome 3 pericentromeric satellites. (A-C) StainedGlass self-identity heatmaps (top) and repeat annotations (middle) of the chromosome 3 pericentromeric regions in A4 (A), SAM (B), and GUM (C) assemblies. Colored alignments indicate local sequence identity among repeated segments. Repeat tracks show the distribution of TE classes and satellite arrays, with the CENP-A-associated region marked with an asterisk. Observed/expected Hi-C contact maps for A4 and SAM (A and B, bottom) reveal elevated contacts within individual *dodeca* arrays and between arrays on opposite sides of the CENP-A domain. D. Model of centromere organization inferred from sequence and Hi-C data. The retroelement-rich CENP-A–associated region forms the centromeric core, flanked by *dodeca* arrays that form a compact, self-interacting domain. Surrounding *Prodsat* arrays are more variable and exhibit weaker long-range interactions, forming a broader pericentromeric compartment. The i-motif schematic illustrates a potential mechanism for higher-order folding of *dodeca* DNA via intercalated C-rich strands, as described in previous biochemical studies.

In contrast, *Prodsat* shows weaker long-range interactions with itself or with other satellite blocks. Hi-C maps reveal some weak contact between the centromere-proximal region and the small *Prodsat* array located nearest the centromere, but this signal is substantially weaker than the interactions observed among *dodeca*-rich regions (mean contact between *Prodsat-Prodsat* and *dodeca-dodeca* bins 1.76 and 4.05, respectively). The high-identity *Prodsat* regions located toward the distal end of the array show especially limited long-range contact, suggesting that the most homogenized portions of *Prodsat* may not contribute strongly to the dominant Cen3 interaction architecture. *Prodsat* is bound by the Proliferation Disrupter protein, which has been proposed to organize this satellite into higher-order heterochromatic structures required for chromosome condensation (Török et al. 2000). Thus, while *Prodsat* may contribute to broader pericentromeric organization, the strongest and most consistent 3D feature is the interaction among *dodeca*-rich regions flanking the CENP-A-associated hub.

This organization is consistent with prior biochemical studies showing that the *dodeca* satellite can adopt non-B DNA structures *in vitro*. These studies show that *dodeca* repeats can form i-motif structures *in vitro* through the intercalation of two C-rich hairpins, with the antiparallel orientation of the paired hairpins enabling C-C⁺ bond formation across strands (Abad and Villasante 2000; Garavís, Méndez-Lago, et al. 2015). Interestingly, we observe that the C-rich strands of the two *dodeca* arrays flanking the CENP-A domain are oriented on opposite strands in all three genomes (Figs. 4A, B, C), precisely the antiparallel geometry that would allow for the formation of intermolecular i-motif structures between arrays. This strand orientation is consistent with a model in which the two flanking arrays interact through non-B DNA contacts when brought into spatial proximity, as suggested by our Hi-C data.

Together, these results support a model in which the *dodeca* arrays flanking Cen3 form a compact, self-interacting domain that brings satellite-rich chromatin on either side of the CENP-A-associated region into close spatial proximity (Fig. 4D). In this model, the retroelement-rich hub marks the functional centromeric core, *dodeca* defines a conserved architectural domain surrounding that core, and *Prodsat* forms a more variable, expansive pericentromeric domain.

## Discussion

Our single-fly assemblies reveal substantial variation in centromere and pericentromere organization within *D. melanogaster*, while preserving a consistent higher-order structure at the chromosome 3 centromere. This extends earlier work showing that fly centromeres are organized as retroelement-rich islands embedded within large satellite arrays, with CENP-A concentrated primarily over the islands and extending only partially into adjacent satellites (Chang et al. 2019). Here, we resolve this architecture in its full pericentromeric context, most clearly at Cen3, where the centromeric island can be placed within a continuous sequence spanning both chromosome arms. Together with CENP-A and Hi-C profiles, these data support a model in which different repeat classes make related but distinct contributions to centromere organization. The retroelement-rich hub marks the centromere-associated core, whereas surrounding satellite arrays contribute to higher-order chromatin structure. Notably, the spatial context of the CENP-A–associated hub is conserved across strains despite extensive variation in the flanking satellite arrays, indicating that structural features of the centromere are maintained even as the underlying DNA sequence evolves.

These findings provide a potential route toward reconciling the centromere paradox. Classical models invoke epigenetic specification, meiotic drive, and compensatory protein evolution to explain how centromere function is maintained despite rapid DNA turnover (Henikoff et al. 2001; Kursel and Malik 2018). Our data support a complementary view in which selection may act not only on primary DNA sequence but also on higher-order chromatin architecture (Plohl et al. 2014; Kasinathan and Henikoff 2018). Consistent with this, *dodeca*-rich arrays flanking the CENP-A-associated hub show conserved, preferential interactions across strains, whereas the hub itself shows relatively limited interactions with the surrounding satellite-rich chromatin. This spatial separation suggests that satellite arrays may form a distinct chromatin compartment surrounding the centromeric island, potentially preserving a local environment accessible to kinetochore-associated proteins while contributing to higher-order packaging. Although Hi-C does not directly measure accessibility or transcription, this framework provides a testable model for how repeat organization could influence centromere function or strength (Chmátal et al. 2014; Kursel and Malik 2018).

Under this model, centromeric SVs may affect centromere strength not only through changes in repeat abundance but by altering the spatial organization of the centromeric island relative to surrounding satellite domains. Expansions or contractions of *dodeca* and *Prodsat* arrays could modify the size or interaction capacity of the satellite-rich compartment, whereas copy number variation in other repeats, such as the Cen3 IGS units, may influence the transcriptional potential of the CENP-A-associated island. These IGS sequences are likely derived from rDNA spacers (Chang et al. 2019), and canonical rDNA IGS units contain transcriptional promoters (Grimaldi and Di Nocera 1988). As transcription of *G2/Jockey-3* elements has been proposed to help maintain the CENP-A chromatin landscape (Chabot et al. 2024), the IGS copy number could influence local transcription, CENP-A maintenance, or kinetochore assembly. SVs that alter these properties could therefore provide a substrate for centromere drive, especially if they generate differences in centromere strength between chromatids during meiosis.

One possible molecular basis for the satellite interactions observed here is the structural potential of satellite DNA. Non-canonical DNA structures, specifically i-motifs, have been proposed or observed in centromeric satellites of *Drosophila* and humans (Abad and Villasante 2000; Garavís, Escaja, et al. 2015; Garavís, Méndez-Lago, et al. 2015; Zeraati et al. 2018; Patchigolla and Mellone 2022). Such folding could contribute to the high-density interactions observed around Cen3 and generate a compact, satellite-rich chromatin environment that surrounds the CENP-A-associated hub but is spatially distinct from it. Although our Hi-C data do not directly demonstrate these structures *in vivo*, they are consistent with a model in which rapidly evolving satellite sequences retain structural properties that support conserved higher-order organization.

More broadly, our data provide a framework for connecting centromeric structural variation, transcriptional potential, and three-dimensional chromatin organization within the TE island–satellite ‘sea’ of *Drosophila* centromeres. The conserved organization described here is based on Hi-C data from adult females and may not fully reflect the centromere conformation in the embryo, germline, or meiotic contexts, where CENP-A establishment and centromere drive are most active, and where interactions between the centromeric region and surrounding satellite arrays may be more extensive. Future work integrating stage-specific chromatin-contact maps with direct measurements of centromeric transcription, satellite array size, IGS copy number, and chromosome segregation will be needed to determine whether these structural features modulate centromere strength. The apparent uncoupling of rapid sequence evolution from conserved spatial organization nevertheless parallels observations at human centromeres, where extensive alpha-satellite variation coexists with preserved three-dimensional architecture (Altemose et al. 2022; Sen Gupta et al. 2023; Gao et al. 2024; Logsdon et al. 2024), suggesting that higher-order chromatin organization may represent a conserved principle of metazoan centromere biology.

## Materials and Methods

### DNA extraction and sequencing

Single-fly DNA was extracted from individual adult female flies (Supplementary Table 1) using the Qiagen Magattract HMW DNA Extraction Kit. Flies were flash-frozen in liquid nitrogen, ground into a fine powder using a chilled plastic pestle, and incubated in lysis buffer at 56 °C for 16 hours prior to purification. DNA libraries were prepared and sequenced using PacBio Revio SPRQ chemistry on a SMRT Cell 25M at the University of Washington Long Read Sequencing Center.

For Oxford Nanopore sequencing of the A4 strain, high-molecular-weight DNA was extracted from pooled adult females as previously described (Chakraborty et al. 2016). DNA fragments shorter than 40 Kb were removed using the PacBio Short Read Eliminator XL kit. Libraries were end-repaired using the NEBNext Companion Module for ONT Ligation Sequencing Kit v2 (New England Biolabs), followed by adaptor ligation with the ONT Duplex-Enabled Ligation Sequencing Kit V14. Sequencing was performed on an R10.4.1 flow cell using a PromethION Solo device for 72 hours.

### Assembly and scaffolding

PacBio HiFi reads were assembled using hifiasm v0.25.0 (Cheng et al. 2021) with default parameters. The primary hifiasm assemblies were used as the primary contig sets for each strain. For the wild-derived SAM and GUM strains, alternate hifiasm assemblies were retained because they captured alternate haplotypes corresponding to the inversion polymorphisms. Microbial contigs were identified by extracting the first, middle, and last 30 Kb of each contig and aligning these sequences to the NCBI nucleotide database (downloaded on 4-23-2024) using BLASTN. Contigs with microbial matches were removed.

Hi-C libraries were generated from two adult females of the A4 and SAM strains using the Arima High Coverage Hi-C Kit (Arima Genomics Services). Reads were adapter-trimmed with fastp v0.23.4 (Chen et al. 2018) and mapped to the contig-level assemblies using the Juicer pipeline v1.6 (Durand, Shamim, et al. 2016). Contact maps were visualized in Juicebox v1.11.08 (Durand, Robinson, et al. 2016) and used to scaffold contigs into the major chromosome arms. We assigned the large *Prodsat*-rich contig to a chromosome based on the Hi-C interactions between the first and last 50 Kb of the *Prodsat* contig and the first and last 50 Kb of all other scaffolds (Supplementary Table 4). The *Prodsat*-contig end that terminates in AAGAT repeats showed the most contacts with the proximal end of the 2L contig, supporting their physical adjacency. We therefore scaffolded this *Prodsat*-rich contig to the 2L arm contig. Final contact maps were balanced and plotted using HiCExplorer (Wolff et al. 2020). GUM contigs were scaffolded with mscaffolder (Chakraborty et al. 2018) with the A4 single-fly assembly as the reference.

### Assembly quality assessment

Assembly quality was initially evaluated by mapping HiFi reads to contigs using minimap2 v2.29 (Li 2017) and inspecting alignments for potential misassemblies. Misjoins in the SAM and GUM assemblies were confirmed by inspecting read alignment and visualizing the unitig graph in Bandage (Wick et al. 2015), after which they were manually corrected. To assess coverage variation within highly repetitive heterochromatic regions, we mapped reads to each assembly using Winnowmap v2.03 (Jain et al. 2020), as it is optimized for repetitive sequences. The 3L–3R centromere-spanning contig showed reduced coverage across the *dodeca* and *Prodsat* satellite arrays flanking the centromere. To estimate the expected coverage for these repetitive regions, we extracted reads that mapped to the low-coverage interval, generated a k-mer histogram using Jellyfish, and analyzed the profile with GenomeScope. This analysis estimated a local read depth of approximately 33×, which is more consistent with the observed coverage in these regions than the genome-wide estimate of 57× from the full read set (Supplemental Fig. 4).

Assembly consensus accuracy (QV) was estimated by mapping HiFi reads back to each assembly and calling variants with PEPPER-Margin-DeepVariant (Shafin et al. 2021). We treated homozygous non-reference (1/1) variants with FILTER = PASS and Genotype Quality >10 as high-confidence discrepancies between the reads and the assembly (Koren et al. 2017; Solares et al. 2018). The number of such sites was divided by the assembly size (bp) to obtain an estimated error rate, which was converted to a Phred-scaled quality value:

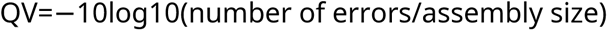

Genome completeness was assessed using BUSCO v5.7.1 (Simão et al. 2015) with the Diptera ortholog database (downloaded on 7-8-24), and assembly contiguity statistics were produced using QUAST (Gurevich et al. 2013).

### Repeat annotation

We annotated repeats in each genome using RepeatMasker v4.1.8 with a *D. melanogaster* repeat library modified from Repbase to include satellite DNA (Khost et al. 2017). The RepeatMasker output file was parsed using a Python script, rm_to_bed.py, to identify simple repeat annotations corresponding to the *Prodsat* sequence or to AATAG, AAGAG, or AAGAT.

### CENPA ChIP-seq analysis

To confirm that the 3L–3R centromere-spanning contig includes the functional centromeric domain, we mapped publicly available CENP-A ChIP-seq data from Oregon-R embryos (Chang et al. 2019) to each contig-level assembly using BWA v0.7.18 (Vasimuddin et al. 2019). Optical duplicates were marked with Picard MarkDuplicates, and alignments were f ltered to retain only mapped, primary alignments with mapping quality ≥30 using SAMtools v1.21 (Danecek et al. 2021). The log2 ratio of CENP-A relative to the input control was then calculated using deepTools v3.5.2 bamCompare (Ramírez et al. 2014), with counts-per-million normalization based on the total number of reads mapped to each assembly. Peaks were called from the Rep2 mapping data using MACS v3.0.3 callpeak (Zhang et al. 2008). We then examined the top 50 peaks to identify contigs that contain or span putative centromeric regions, based on their repeat composition and structural similarity to previously characterized *Drosophila* centromere sequences.

### Alignment of peri/centromeric regions

Highly repetitive sequences comprising pericentric heterochromatin and centromeres were aligned using UniAligner v0.1 (Bzikadze and Pevzner 2023). UniAligner CIGAR output was parsed into non-overlapping 1 kb windows, and percent identity was calculated for each window as:

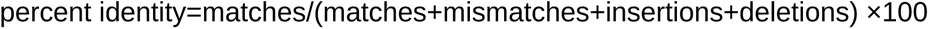

To visualize the structural differences between the retroelement-rich CENP-A islands recovered in our assemblies and the previously characterized iso-1 assembly (Chang et al. 2019), we performed pairwise alignments of iso-1 with A4, A4 with GUM, and GUM with SAM. The alignments were visualized as ribbon plots using the R package SVbyEye (Porubsky et al. 2025), with ribbons colored by percent identity. The self-identity heatmaps were produced using the StainedGlass snakemake pipeline with a 2 Kb window size (Vollger et al. 2022).

### Quantification of contact conservation and satellite-pair Hi-C contacts

To quantify conservation of contact organization between strains, one-to-one corresponding 25 Kb bins between A4 and SAM chromosome 3 pericentromeres were identif ed using UniAligner alignments. Bin pairs were retained if they were reciprocal best matches, shared ≥ 5 Kb of aligned sequence, and contained ≥ 50% aligned sequence. Hi-C contact values were transformed to log2 observed/expected, with expected contact frequencies calculated separately for each genomic distance diagonal using nonzero contacts only. Contact strengths among bin pairs with nonzero contacts in both strains were compared using Pearson and Spearman correlations. For each corresponding bin, we also calculated its mean observed/expected interaction strength across all detected contacts and compared these bin-level interaction profiles between strains. To test whether the observed similarity exceeded what would be expected by chance, we circularly shifted the SAM bin correspondence relative to A4 while preserving the order and internal structure of each contact map. The correlation was recalculated for every possible nonzero shift, and the *p*-value was defined as the fraction of shifted comparisons with a correlation equal to or greater than the observed value.

To quantify interactions among satellite classes, we divided the pericentromeric interval of chromosome 3 into 25-Kb bins matching the Hi-C matrix resolution. Bins were intersected with RepeatMasker annotations and classified by dominant repeat content (>=50%) as *dodeca*-rich or *Prodsat*-rich. Hi-C contact counts were modeled separately for A4 and SAM using negative binomial mixed-effect models in glmmTMB. Satellite-pair class was included as a fixed effect, log-transformed genomic distance as a covariate, and the distance bin and identities of the two interacting bins were included as random effects. Pairwise contrasts among satellite-pair classes were estimated with Benjamini–Hochberg correction for multiple testing. To ensure that enrichment estimates were not driven by zero-contact bin pairs, the analysis was repeated after excluding such pairs.

## Data Access

All sequencing reads and genome assemblies generated in this study have been submitted to the NCBI BioProject database (https://www.ncbi.nlm.nih.gov/bioproject/) under accession Number PRJNA1482540. All scripts for genome assembly and analysis are available as a Supplemental Code file and on GitHub at https://github.com/chakrabortymlab/cenvar.

## Acknowledgements

We sincerely thank Dr. Rebecca Yang for providing the GUM and SAM stocks. We also thank Adolfo A. Delgado and Harsh G. Shukla for valuable discussions on repetitive sequence analysis, and Trevor Millar for advice on statistical analyses. This work was supported by the Texas A&M startup funds and the NIH grant R00GM129411 to M.C.

