## Supplemental Materials for "A conserved architectural domain shapes centromere evolution in *Drosophila*"

Affiliations:
Department of Biology, Texas A&M University, College Station, TX 77843

### Supplemental Figures

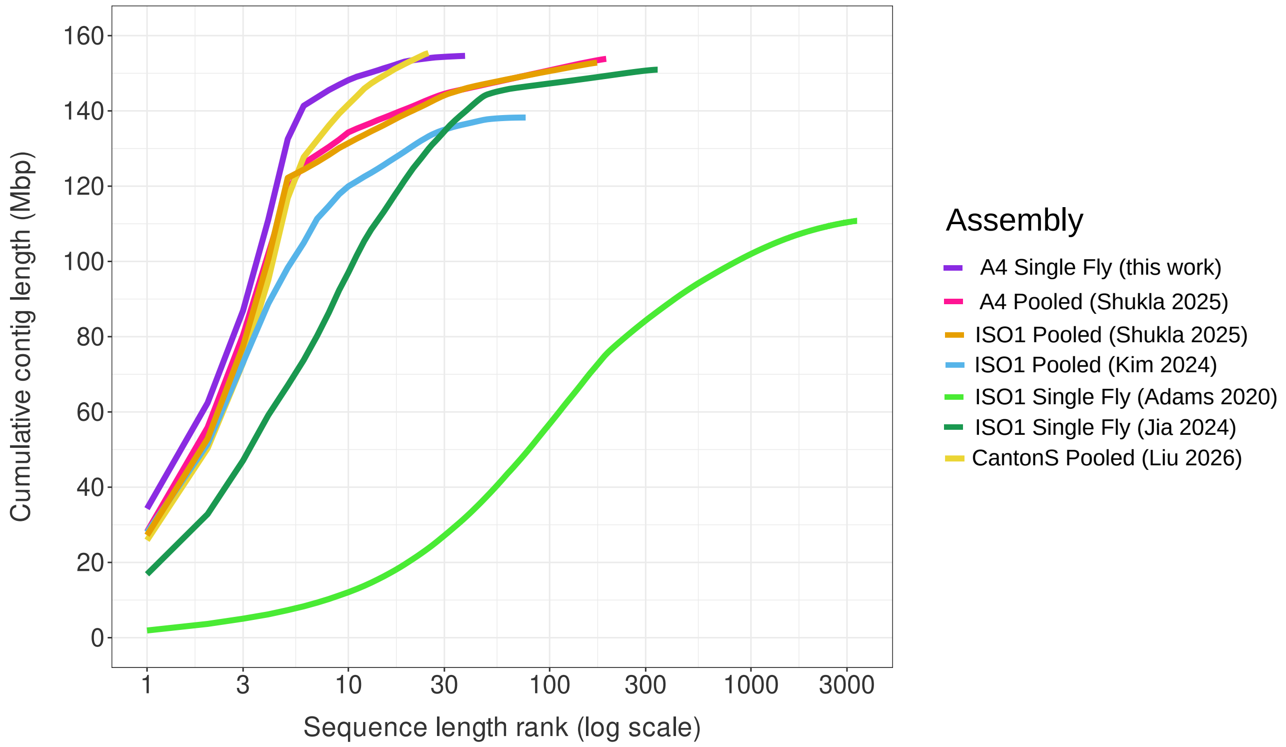


**Supplemental Figure 1.** Comparison of contiguity among *Drosophila melanogaster* genome assemblies. Cumulative assembled sequence length is plotted against contig rank (log scale) for previously published assemblies generated from either single flies or pooled samples (Adams et al. 2020; Jia et al. 2024; Kim et al. 2024; Shukla et al. 2025), together with the single-fly A4 assembly generated in this study. Scaffolded assemblies were split at gap sequences (runs of Ns) to enable contig-level comparisons. Putative Y-linked contigs were excluded from other assemblies because our A4 assembly was generated from a female.

.

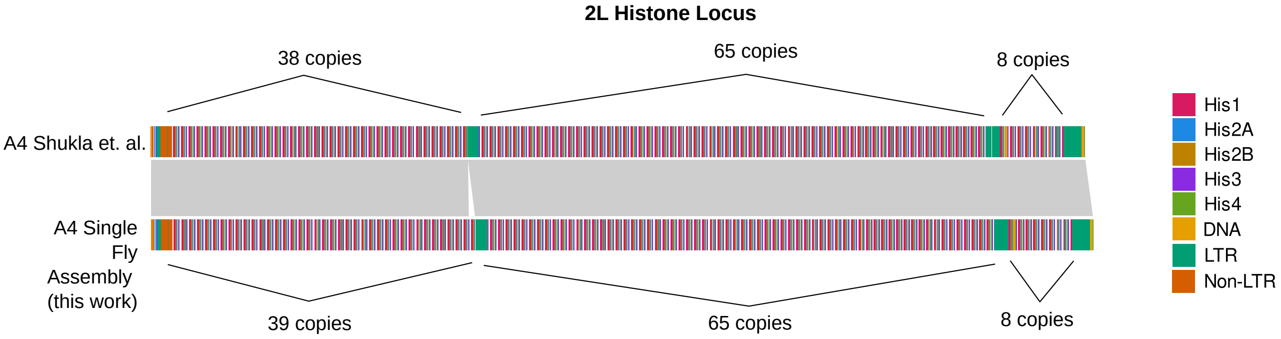


**Supplemental Figure 2.** Single-fly assembly resolves the chromosome 2L histone gene cluster. Comparison of the 2L histone locus in the A4 single-fly assembly generated here and a previously published A4 assembly (Shukla et al. 2025), where a targeted assembly approach was used to resolve this region. Colored blocks denote histone genes and interspersed repeat classes; grey ribbons indicate homologous sequence between assemblies. The single-copy difference in the first block may reflect biological variation among A4 samples or rapid turnover within the cluster.

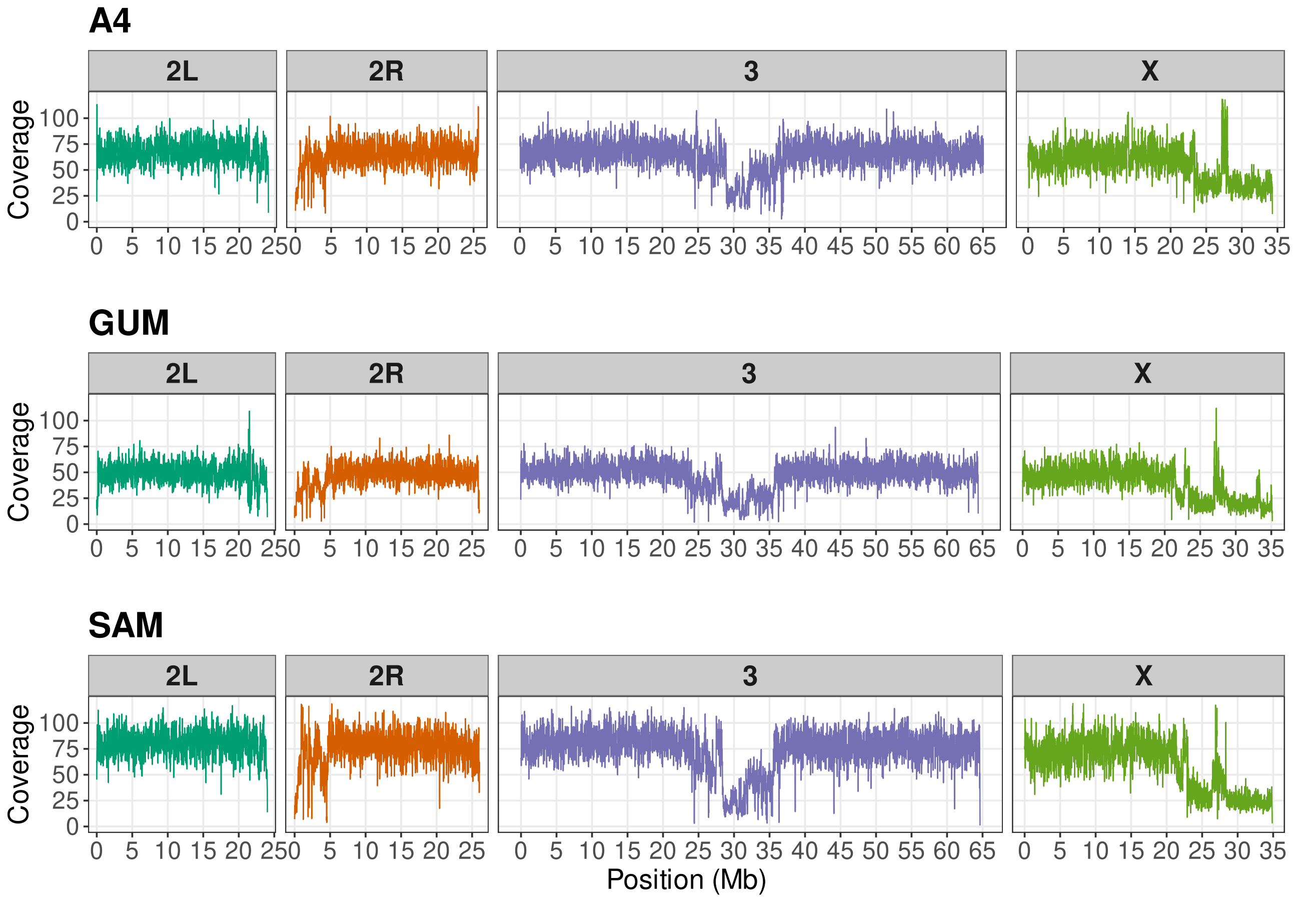


**Supplemental Figure 3.** Sequencing coverage across major chromosome scaffolds in the A4, GUM, and SAM single-fly assemblies. Coverage profiles are shown along the 2L, 2R, chromosome 3, and X chromosome scaffolds for each assembly. Coverage is broadly consistent across euchromatic portions of the chromosome arms. In contrast, pericentric heterochromatin shows lower and more uneven coverage, as expected for highly repetitive heterochromatic regions (Nurk et al. 2022; Xie et al. 2025)

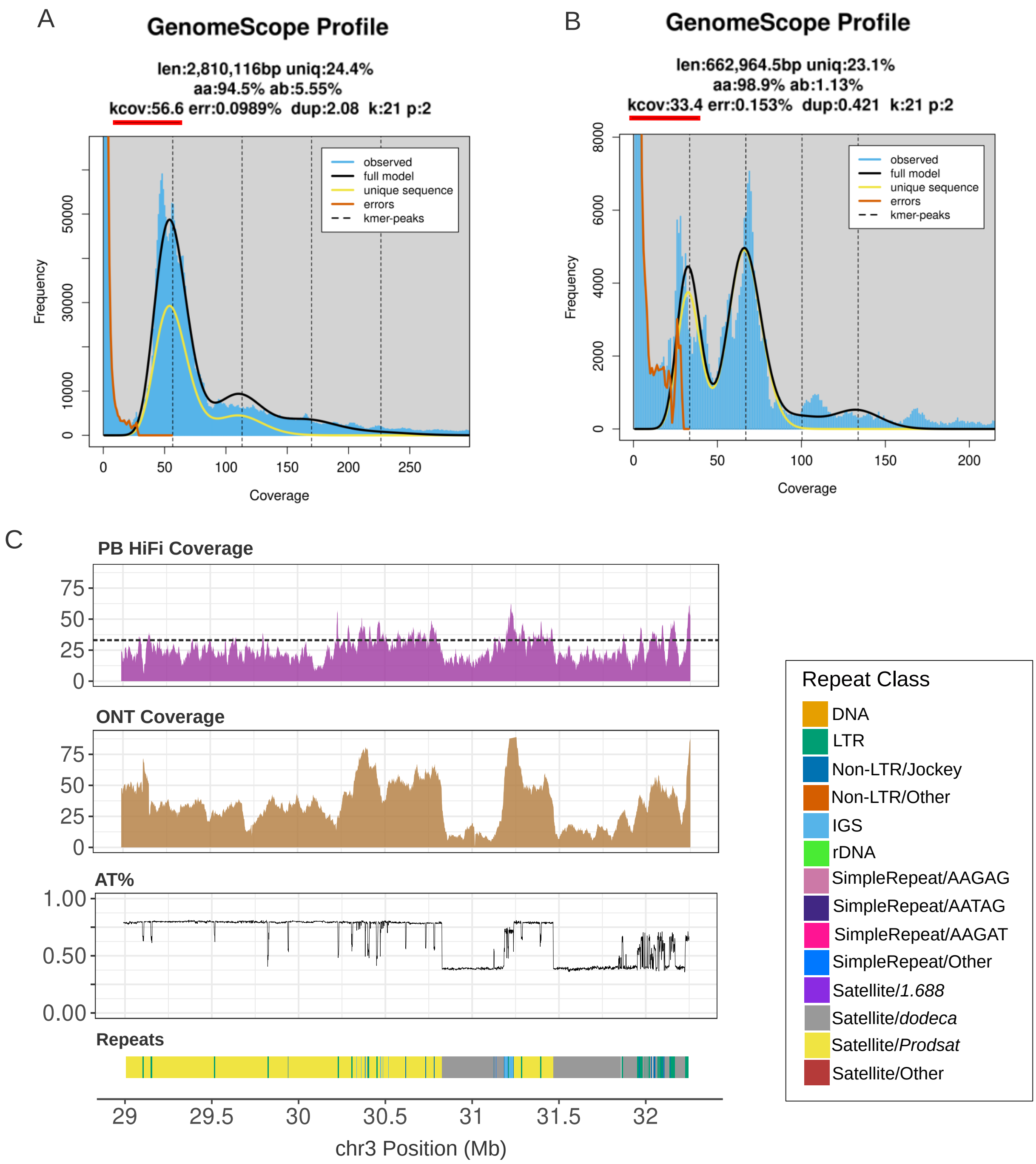


**Supplemental Figure 4.** Reduced sequencing coverage across the pericentromeric region of chromosome 3. (A) GenomeScope k-mer profile of PacBio HiFi reads from a single A4 female, estimating a genome-wide coverage of approximately 57×. (B) GenomeScope profile generated from reads mapping to the chromosome 3 pericentromeric interval, including centromere-proximal satellite arrays, which estimates lower coverage (approximately 33.4×). (C) PacBio HiFi coverage from the single A4 female and Oxford Nanopore coverage from pooled A4. While PacBio HiFi coverage is reduced across the satellite-rich sequences, it is consistent with the lower expectation indicated by the GenomeScope profile for this region (black dotted line).

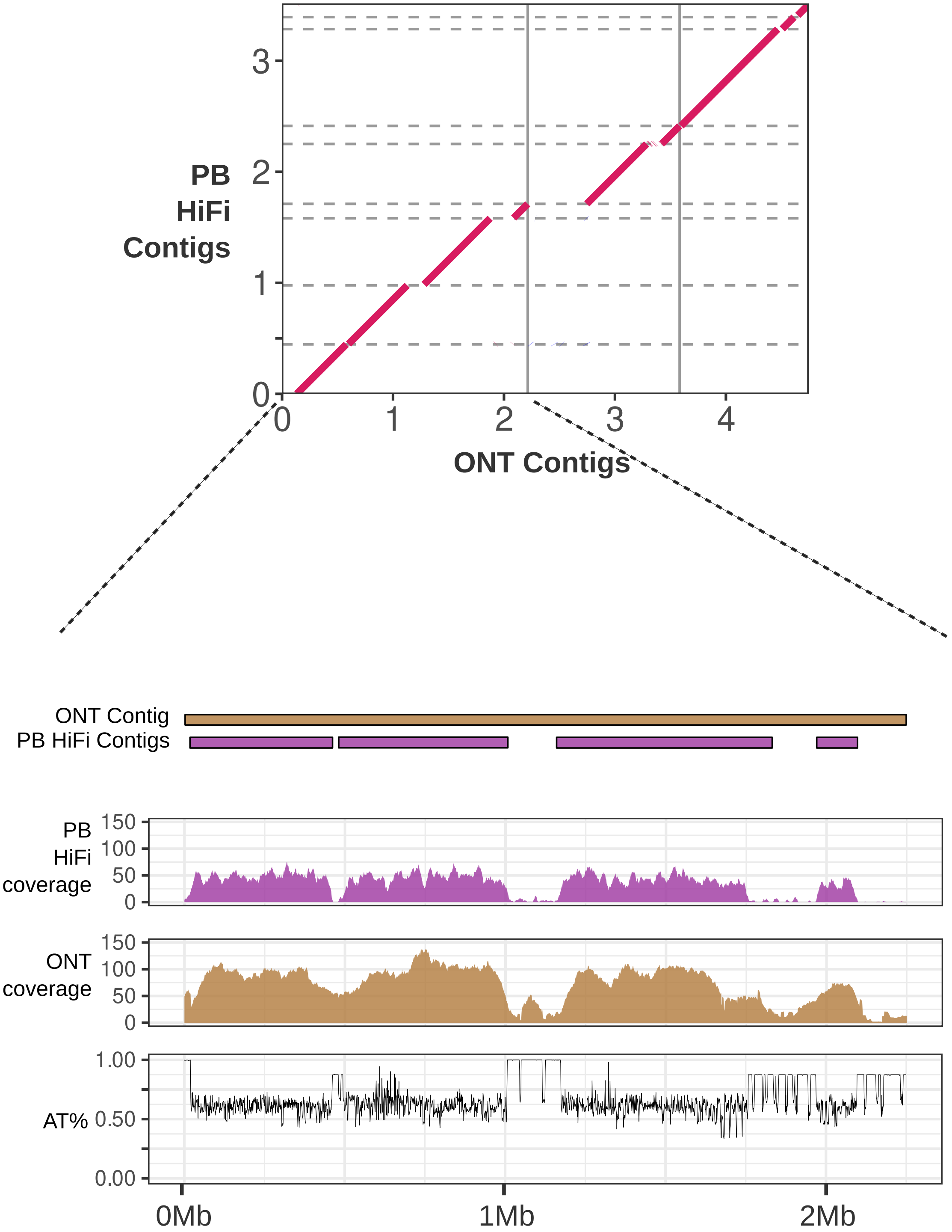


**Supplemental Figure 5.** Fragmentation of an AT-rich 3R pericentromeric region in the PacBio HiFi assembly. An ONT contig spanning this region corresponds to four PacBio HiFi contigs, as illustrated by the dot plot and the alignment schematic. PacBio HiFi coverage from a single A4 female and ONT coverage from pooled A4 females are shown across the region, together with local AT content. Breaks between HiFi contigs (pink) coincide with intervals of reduced HiFi coverage and elevated AT content, consistent with reduced recovery of highly AT-rich sequence in the HiFi-only assembly (Shukla et al. 2025; Carvalho et al. 2026).

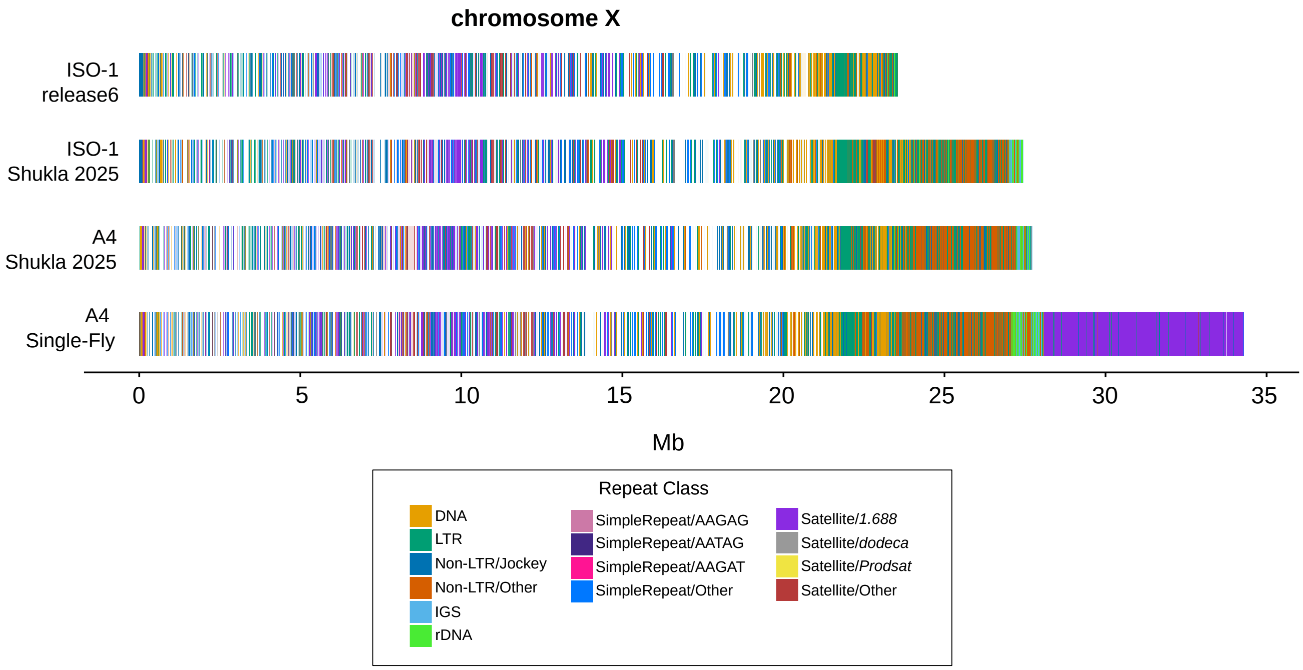


**Supplemental Figure 6.** Comparison of repeat organization across the X chromosome in iso-1 release 6, the iso-1 and A4 assemblies of (Shukla et al. 2025), and the A4 single-fly assembly generated in this study. Whereas the iso-1 release 6 X chromosome assembly lacks much of the heterochromatic sequence proximal to the centromere, the HiFi-based iso-1 and A4 assemblies of (Shukla et al. 2025) extend partially into the X-linked rDNA array. Our single-fly assembly further recovers the rDNA array and an approximately 6-Mb 1.688 satellite array, all within a single contig, which terminates in simple 5-bp repeats previously inferred to lie adjacent to the X centromere.

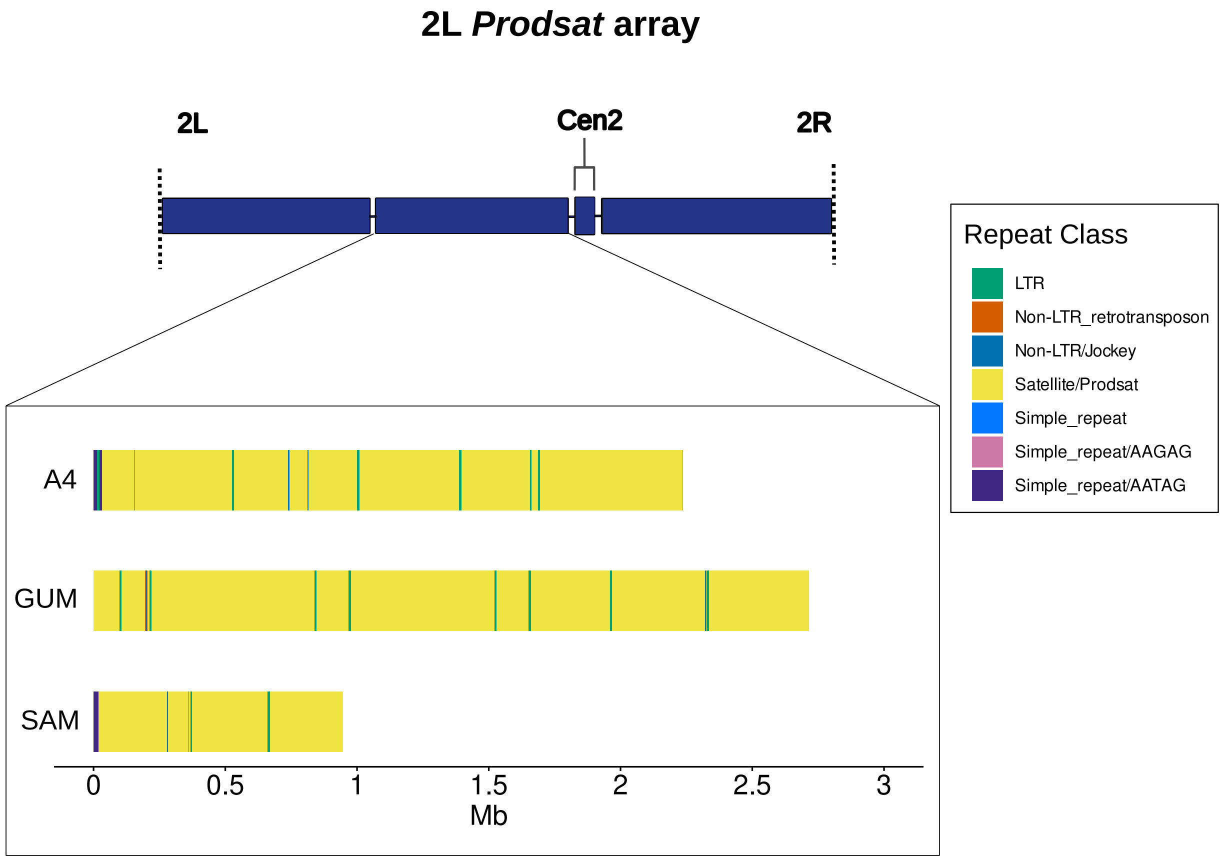


**Supplemental Figure 7.** Schematic placement of the 2L *Prodsat* contig relative to 2L, Cen2, and 2R. This contig was positioned between 2L and Cen2 based on its expected cytological location and Hi-C contacts with adjacent chromosome 2 sequences. Bottom, repeat annotations of the corresponding contigs in A4, GUM, and SAM. In all three assemblies, the contig is composed predominantly of *Prodsat* satellite sequence interspersed with LTR retrotransposons and bounded by simple 5-bp repeats. The overall array length and internal organization differ among strains, consistent with rapid turnover of *Prodsat* arrays.

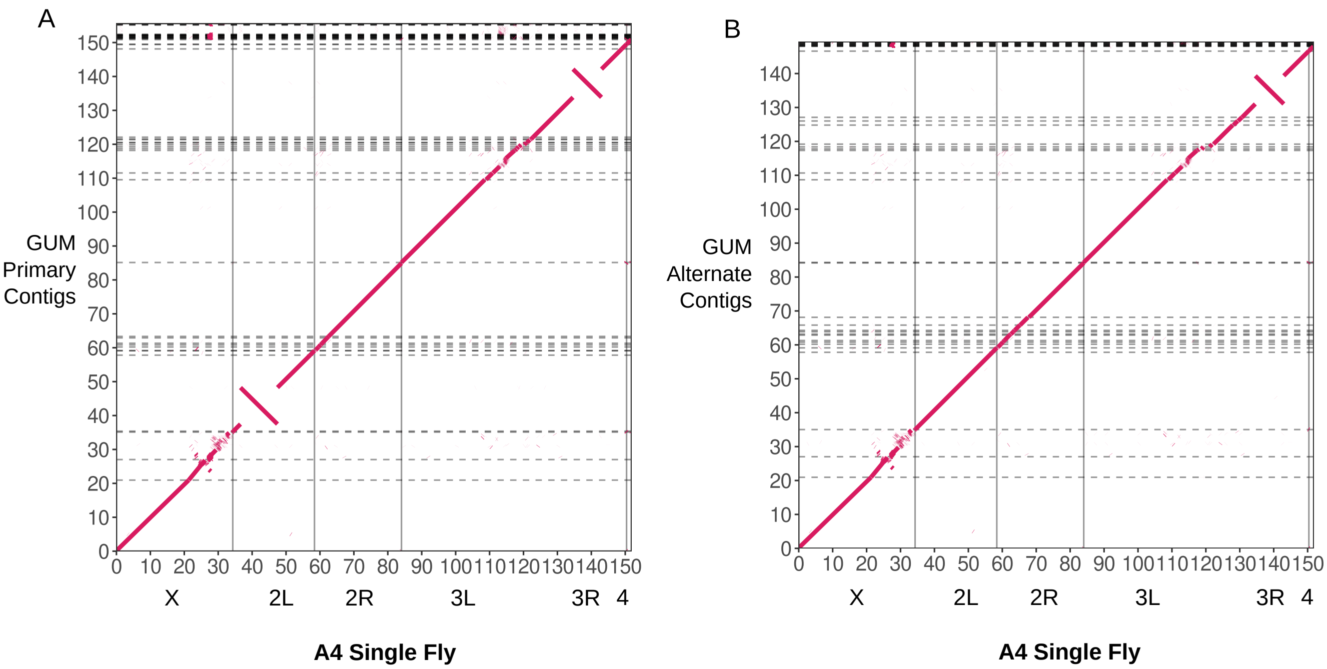


**Supplemental Figure 8.** Dot plots showing comparisons between the GUM primary (A) and alternate (B) contig sets with the scaffolded chromosome arms of the A4 single-fly assembly. Both GUM assemblies contain a contig that spans the chromosome 3 centromere and joins the 3L and 3R arms. The two assemblies recover the standard and inverted arrangements of *In(2L)t*, whereas only the inverted arrangement of *In(3R)Payne* is observed.

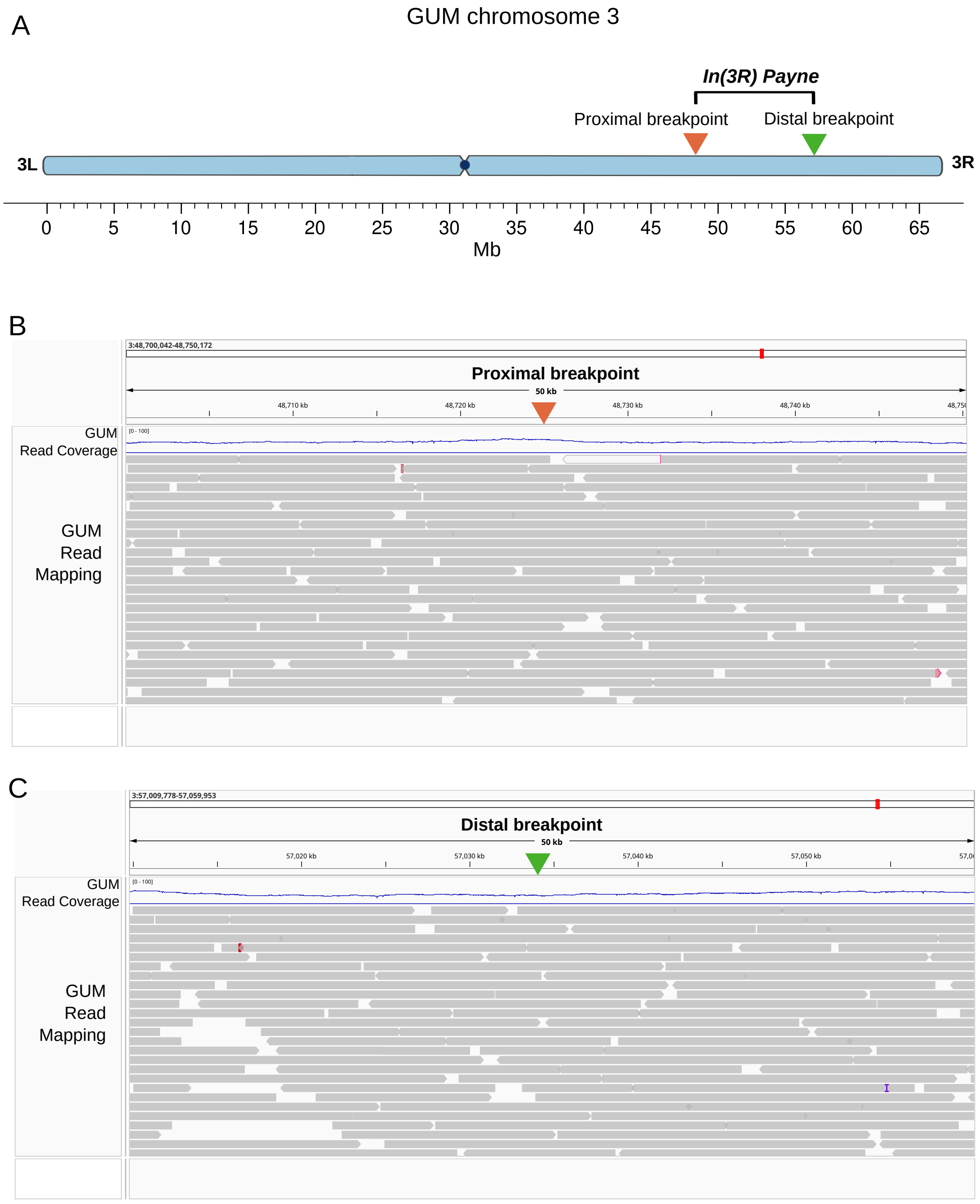


**Supplemental Figure 9.** GUM is homozygous for the inverted *In(3R)Payne* arrangement. (A) Positions of the proximal and distal *In(3R)Payne* breakpoints on GUM chromosome 3. (B, C) PacBio HiFi reads from the GUM single-fly library mapped to the GUM primary assembly, which carries the inverted arrangement, at the proximal (B) and distal (C) breakpoint junctions. Uniform coverage across both junctions and the absence of split reads, which would suggest the standard arrangement, indicate that the sequenced GUM individual is homozygous for the inversion.

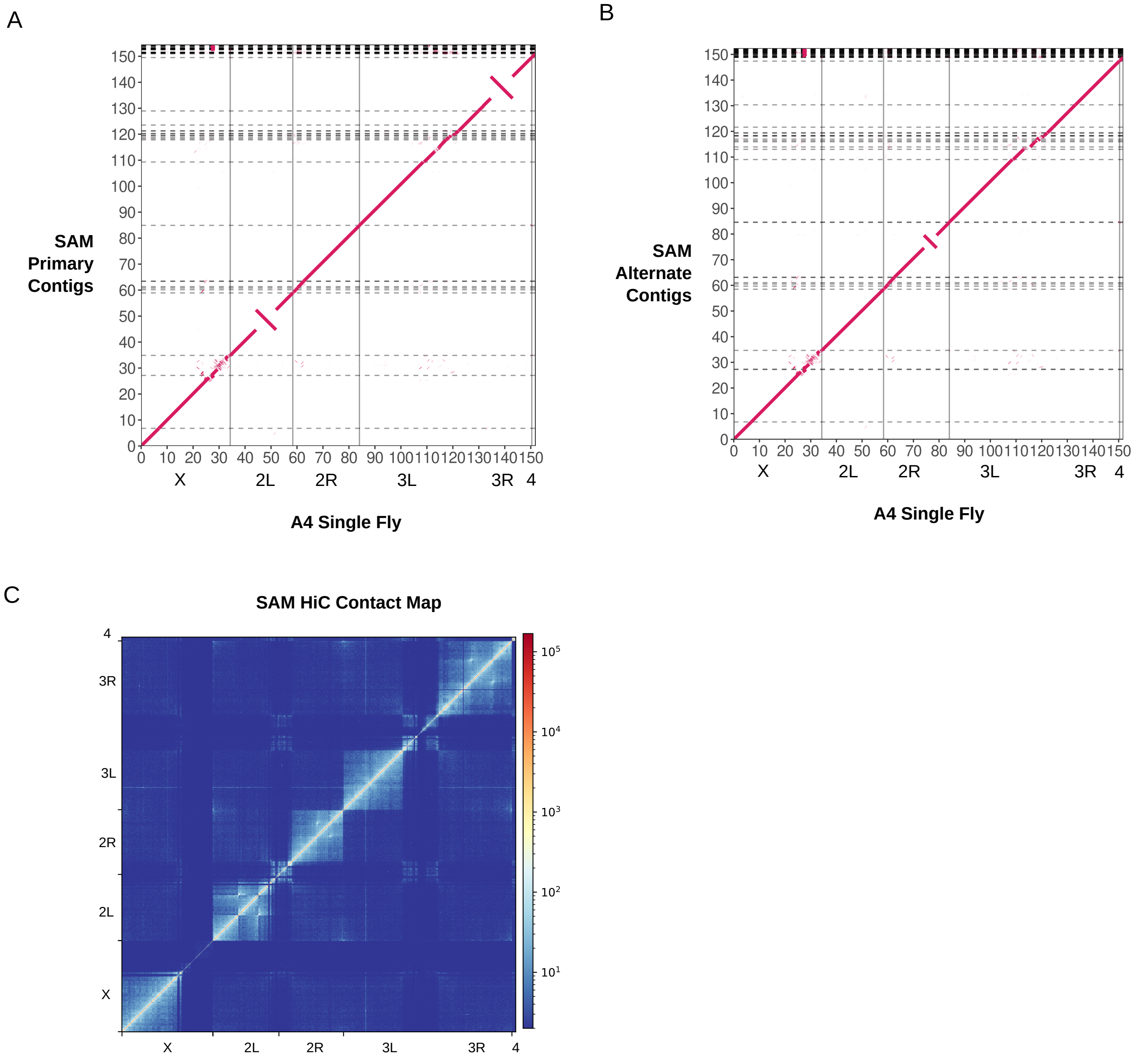


**Supplemental Figure 10.** Dot plots comparing the SAM primary (A) and alternate (B) contigs with the scaffolded A4 single-fly assembly. The primary assembly contains a contig that spans the chromosome 3 centromere and joins the 3L and 3R arms. Together, the two assemblies recover inverted and standard arrangements of *In(3R)Payne* and *In(2R)NS*, as well as a previously undocumented inversion on chromosome 2L. (C) Chromosome-scale Hi-C contact map for the SAM primary assembly. The inversion polymorphisms in 2L, 2R, and 3R are visible as off-diagonal high-density contacts, hallmarks of inverted sequences.

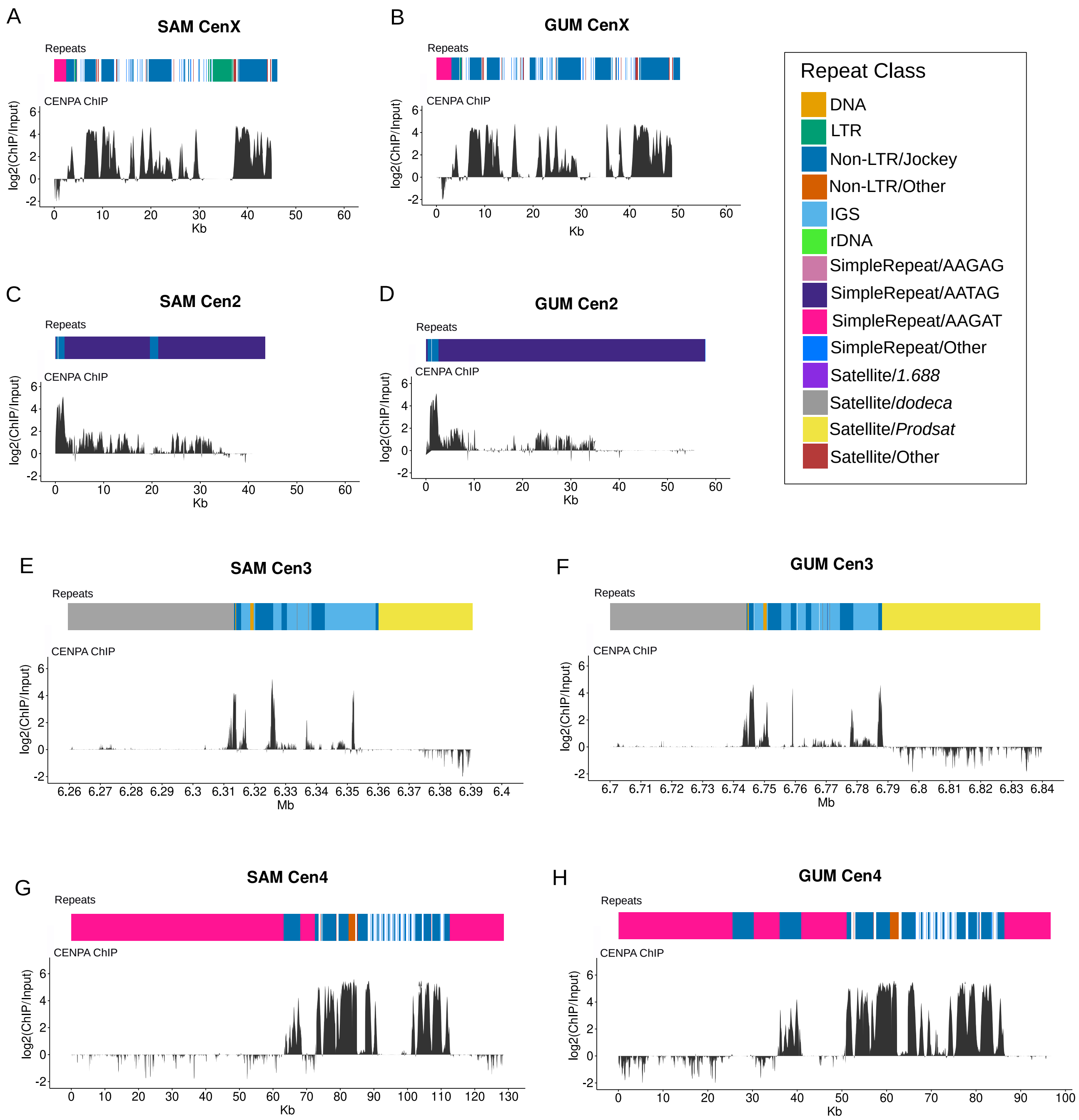


**Supplemental Figure 11.** CENP-A enrichment across centromeric contigs in the SAM and GUM assemblies. CENP-A ChIP-seq reads from OreR embryos (Chang et al. 2019) were mapped to the SAM and GUM single-fly assemblies. Tracks show $\log_2$(ChIP/input) enrichment across CenX (A, B), Cen2 (C, D), Cen3 (E, F), and Cen4 (G, H), with repeat annotations shown above each profile. CENP-A enrichment localizes to TE-rich regions of the assembled centromeric contigs.

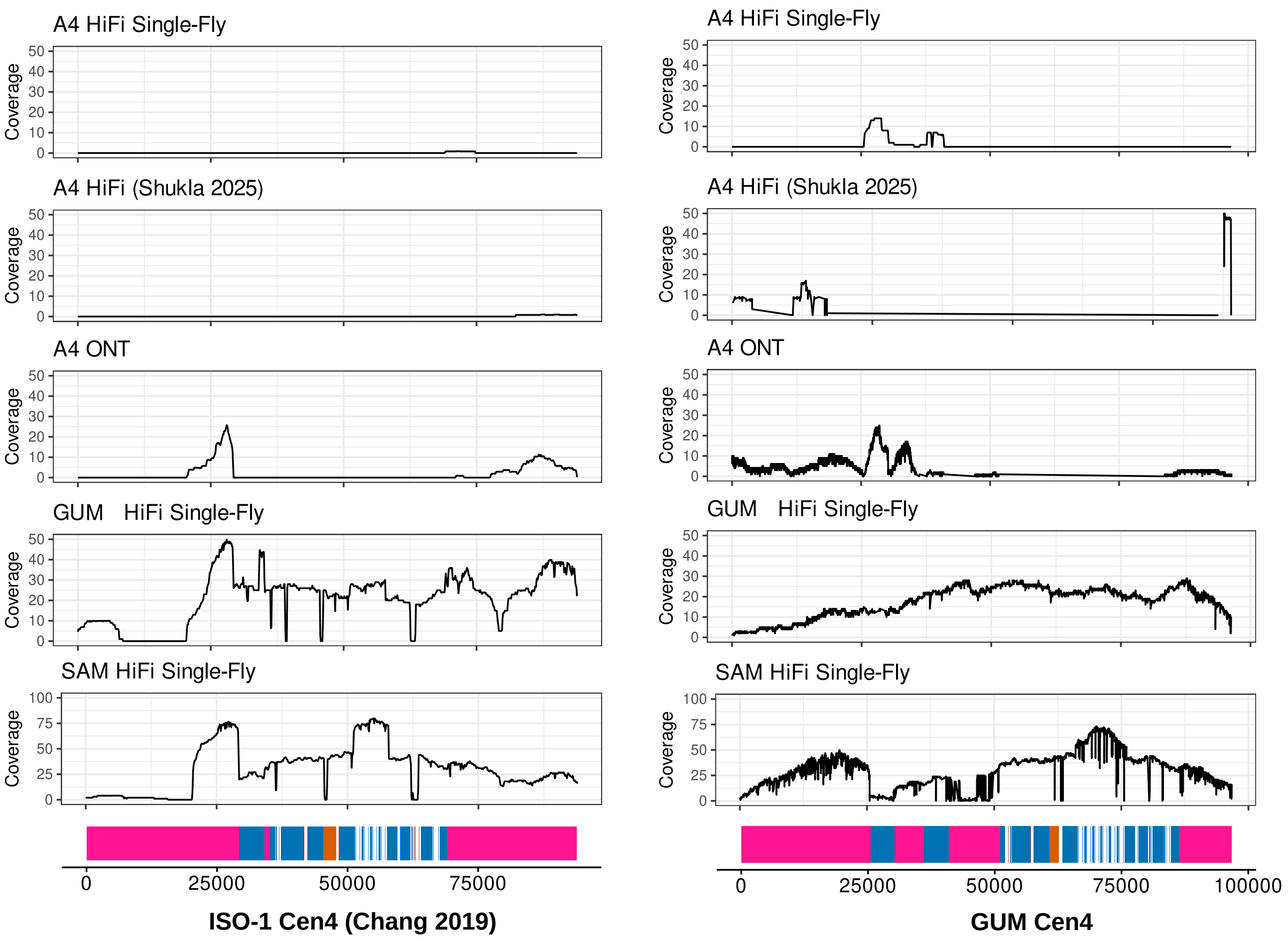


**Supplemental Figure 12.** The chromosome 4 centromeric sequence described for iso-1 is absent or highly diverged in A4. Read coverage from multiple A4 datasets, including single-fly PacBio HiFi, previously published A4 HiFi (Shukla et al. 2025), and pooled A4 ONT reads, was examined across the iso-1 Cen4 sequence characterized by (Chang et al. 2019)(left) and the Cen4 contig recovered in the GUM assembly (right). No corresponding Cen4 sequence was recovered in the A4 single-fly assembly or other A4 assemblies, and A4 reads show little or no coverage across most of either Cen4 sequence. In contrast, GUM and SAM HiFi reads support both the iso-1 and GUM Cen4 sequences.

**Supplemental Figure 13.** Pairwise sequence divergence across chromosome 3. Divergence was calculated in 50-kb windows between the A4 chromosome 3 scaffold and the corresponding GUM (top) and SAM (bottom) scaffolds. Divergence was calculated for each window as:
(#mismatches + #insertions + #deletions) / (#matches + #mismatches + #insertions +#deletions). The schematic below indicates euchromatic, heterochromatic, and centromeric intervals, including the *Prodsat* and *dodeca*-rich region around the centromere. Both comparisons show localized increases in divergence near the centromeric/pericentromeric region relative to most chromosome-arm windows.

### Supplemental Tables

**Supplemental Table 1.** Fly geographic origin and sequencing details

| Strain | Origin | HiFi Reads | Read N50 | Median QV | Yield (Gbp) |
|---|---|---|---|---|---|
| A4 | Zimbabwe, DSPR founder line | 653,594 | 16,369 | 38 | 9.62 |
| GUM | Guam, Oceania. DSSC stock ID: 14021-0231.198 | 512,339 | 15,841 | 38 | 7.14 |
| SAM | American Samoa: DSSC stock ID: 14021-0231.134 | 788,432 | 16,727 | 38 | 11.20 |

| Assembly | Contigs | | | Scaffolds | | Diptera Complete BUSCOs |
|---|---|---|---|---|---|---|
| | Length (bp) | # | N50 (bp) | # | N50 (bp) | |
| GUM alternate | 149,614,805 | 45 | 19,522,920 | 24 | 35,027,080 | 98.5% |
| SAM alternate | 152,748,555 | 78 | 20,418,462 | 57 | 34,524,392 | 99.3% |

**Supplemental Table 2.** GUM and SAM alternate assembly stats

**Supplemental Table 3.** GUM and SAM inversions

| Strain | Assembly | Reference Coordinates (iso-1 r6) | Query Coordinates | Known as |
|---|---|---|---|---|
| GUM | primary | 2L:2,225,672-13,154,047 | 2L:2,445,749-13,135,749 | In(2L)t, FBab0004696 |
| | primary | 3R:16,428,301-24,751,017 | 3:48,724,950-57,034,014 | In(3R)Payne, FBab0005639 |
| SAM | primary | 2L:9,852,450-17,610,343 | 2L:9,906,922-17,522,754 | NA |
| | primary | 3R:16,428,301-24,751,017 | 3:48,927,574-57,324,300 | In(3R)Payne, FBab0005639 |
| | alternate | 2R:15,389,072-20,276,397 | 2R:16,086,180-21,025,154 | In(2R)NS, FBab0005032 |

**Supplemental Table 4.** A4 *Prodsat* contig contacts with major scaffolds

| Query | QueryStart:TargetStart | QueryStart:TargetEnd | QueryEnd:TargetStart | QueryEnd:TargetEnd |
|---|---|---|---|---|
| X | 1 | 0 | 1 | 0 |
| cenX | 1 | 0 | 1 | 0 |
| 2L | 3 | 0 | 32 | 9 |
| cen2 | 6 | 2 | 6 | 2 |
| 2R | 9 | 6 | 2 | 0 |
| 3L_3R | 2 | 0 | 2 | 0 |
| 4 | 3 | 2 | 3 | 2 |
